## Supplementary Information for "Structure of the membrane-bound formate hydrogenlyase complex from *Escherichia coli*"

**The PDF file includes:**

Supplementary Table S1 to S3

Supplementary Figures S1 to S14

References

**Table S1 Cryo-EM data collection, refinement and model validation statistics.**

| | FHL anaerobic | FHL aerobic |
|---|---|---|
| ***Map EMDB ID*** | **EMD-14429** | **EMD-14430** |
| **Data collection** | | |
| Microscope | Titan Krios G2 | Titan Krios G2 |
| Camera | Gatan K3 Bioquantum | Gatan K2 Summit |
| Voltage (kV) | 300 | 300 |
| Nominal magnification | 105,000x | 165,000x |
| Calibrated pixel size (Å) | 0.831* | 0.828 |
| Dose ($e^-$/Å$^2$) | 61 | 72 |
| Number of frames per image | 60 | 40 |
| Defocus range (µm) | -2.5 to -1.0 | -2.2 to -1.6 |
| **Image processing** | | |
| Motion correction software | MotionCor2 | MotionCor2 |
| CTF estimation software | CTFFIND4 | CTFFIND4 |
| Particle selection software | Topaz | crYOLO |
| Final micrographs (no.) | 7,338 | 1,185 |
| Initial particle images (no.) | 686,602 | 200,409 |
| Final particle images (no.) | 300,386 | 90,459 |
| Map sharpening B-Factor (Å$^2$) | -66.6 | -** |
| Final resolution (Å) | 2.6 | 3.0 – 3.4** |
| ***Model PDB ID*** | **7Z0S** | **7Z0T** |
| **Refinement** | | |
| Modeling software | Coot, Phenix | Coot, Phenix |
| Protein residues | 2,026 | 2,690 |
| Water | 121 | - |
| Ligands | | |
| NI | 1 | 1 |
| 6MO | - | 1 |
| FE | 1 | 1 |
| MGD | - | 2 |
| SEC | - | 1 |
| SF4 | 7 | 8 |
| FCO | 1 | 1 |
| DR9 | 1 | - |
| PTY | 2 | - |
| CDL | 1 | - |
| LMN | 1 | - |
| **Validation** | | |
| MolProbity score | 1.38 | 1.95 |
| Clash score | 4.96 | 8.79 |
| Ramachandran plot (%) | | |
| Outliers | 0.05 | 0.30 |
| Allowed | 2.58 | 4.42 |
| Favored | 97.37 | 95.28 |
| Rotamer outliers (%) | 0.91 | 1.55 |
| Cβ outliers (%) | 0.27 | 1.46 |
| Peptide plane (%) | | |
| Cis proline/general | 5.3/0.0 | 5.0/0.0 |
| Twisted proline/general | 0.0/0.0 | 0.0/0.0 |
| CaBLAM outliers (%) | 1.15 | 2.64 |

* During refinement a pixel size of 0.837 Å was used. The final map was postprocessed using the calibrated pixel size of 0.831 Å.
** EMD-14430 is a composite map. The consensus and focused maps that contributed to the composite map are listed in Table S2.

**Table S2 Cryo-EM data collection and refinement of maps that contributed to the composite map EMD-14430.**

| | EMD-14431 | EMD-14432 | EMD-14433 | EMD-14434 |
|---|---|---|---|---|
| **Data collection** | | | | |
| Microscope | Titan Krios G2 | Titan Krios G2 | Titan Krios G2 | Titan Krios G2 |
| Camera | Gatan K2 Summit | Gatan K2 Summit | Gatan K2 Summit | Gatan K2 Summit |
| Voltage (kV) | 300 | 300 | 300 | 300 |
| Nominal magnification | 165,000x | 165,000x | 165,000x | 165,000x |
| Calibrated pixel size (Å) | 0.828 | 0.828 | 0.828 | 0.828 |
| Dose ($e^-$/Å$^2$) | 72 | 72 | 72 | 72 |
| Number of frames per image | 40 | 40 | 40 | 40 |
| Defocus range (μm) | -2.2 to -1.6 | -2.2 to -1.6 | -2.2 to -1.6 | -2.2 to -1.6 |
| **Image processing** | | | | |
| Motion correction software | MotionCor2 | MotionCor2 | MotionCor2 | MotionCor2 |
| CTF estimation software | CTFFIND4 | CTFFIND4 | CTFFIND4 | CTFFIND4 |
| Particle selection software | crYOLO | crYOLO | crYOLO | crYOLO |
| Final micrographs (no.) | 1,185 | 1,185 | 1,185 | 1,185 |
| Initial particle images (no.) | 200,409 | 200,409 | 200,409 | 200,409 |
| Final particle images (no.) | 90,459 | 90,459 | 90,459 | 90,459 |
| Map sharpening B-Factor (Å$^2$) | -54.5 | -48.0 | -43.7 | -74.2 |
| Final resolution (Å) | 3.4 | 3.1 | 3.0 | 3.4 |

**Table S3 Homologous subunits in complex I, FHL, MBH and soluble [NiFe] hydrogenases.**

| | **Complex I**<br>*H. sapiens /*<br>*Y. lipolytica* | <br><br>*T. thermophilus* | **FHL**<br><br>*E. coli* | **MBH**<br><br>*P. furiosus* | **[NiFe] hydrogenase**<br><br>*D. vulgaris* |
|---|---|---|---|---|---|
| **substrate oxidation** | NDUFS1 | Nqo3 | FdhF (C-term)<br>HycB (N-term) | | |
| | NDUFV1 | Nqo1 | | | |
| | NDUFV2 | Nqo2 | | | |
| **substrate reduction** | NDUFS2 | Nqo4 | HycE (C-term) | MbhL | HydB |
| | NDUFS3 | Nqo5 | HycE (N-term) | MbhK | |
| | NDUFS7 | Nqo6 | HycG | MbhJ | HydA |
| | NDUFS8 | Nqo9 | HycF | MbhN | |
| **(putative) proton trans-location** | ND1 | Nqo8 | HycD | MbhM | |
| | ND2/ND4/ND5 | Nqo14/Nqo13/Nqo12 | HycC (N-term) | MbhH | |
| | ND3 | Nqo7 | HycC (C-term) | MbhI (N-term) | |
| | ND6 | Nqo10 | | MbhD/MbhE | |
| | ND4L | Nqo11 | | MbhG | |
| **sodium trans-location** | | | | MbhF | |
| | | | | MbhA | |
| | | | | MbhB | |
| | | | | MbhC | |

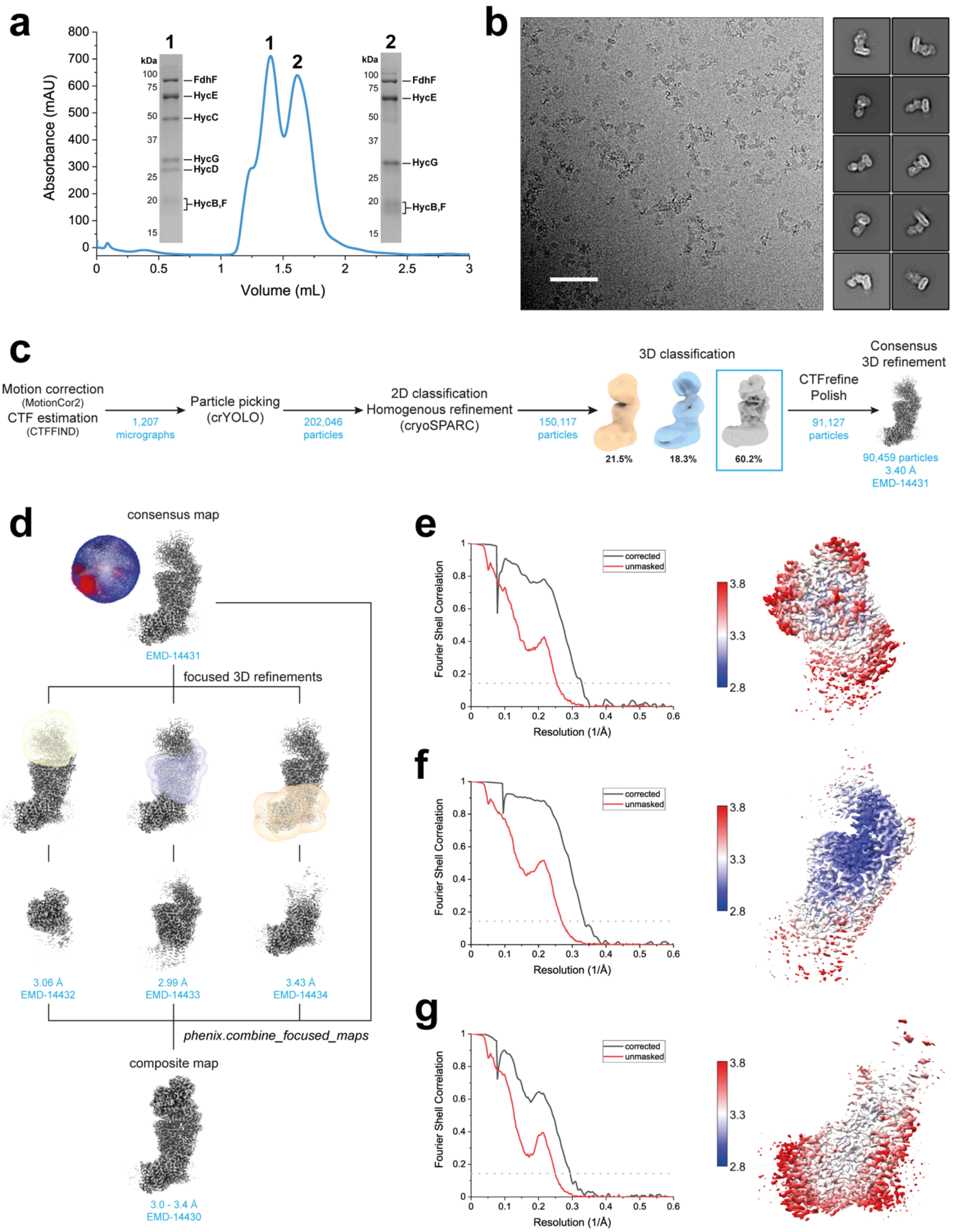


**Fig. S1 Sample preparation and data processing of aerobically prepared FHL.** (**a**) Representative size exclusion chromatography (SEC) profile (Superdex 200 5/150 GL) and SDS-PAGE of peak fractions corresponding to peak 1 (entire heptameric complex) and peak 2 (soluble arm subunits). Gel bands are labeled according to molecular mass. (**b**) Representative cryo-EM micrograph (scale bar 500 Å) and 2D class averages. (**c**) Processing workflow, with all steps performed in RELION-3 except when otherwise indicated. A consensus 3D refinement yields a map (EMD-14431) with a resolution of 3.4 Å. (**d**) The consensus map served as a basis for masked focused 3D refinement, giving focused maps of all regions of the complex (EMD-14432 – EMD-14434) with resolutions between 3.0 - 3.4 Å, which were combined using *phenix.combine_focused_maps*. The resulting composite map (EMD-14430) was used for model refinement. Fourier Shell Correlation (FSC) curve and local resolution estimation for focused refinements EMD-14432 (**e**), EMD-14433 (**f**) and EMD-14434 (**g**).

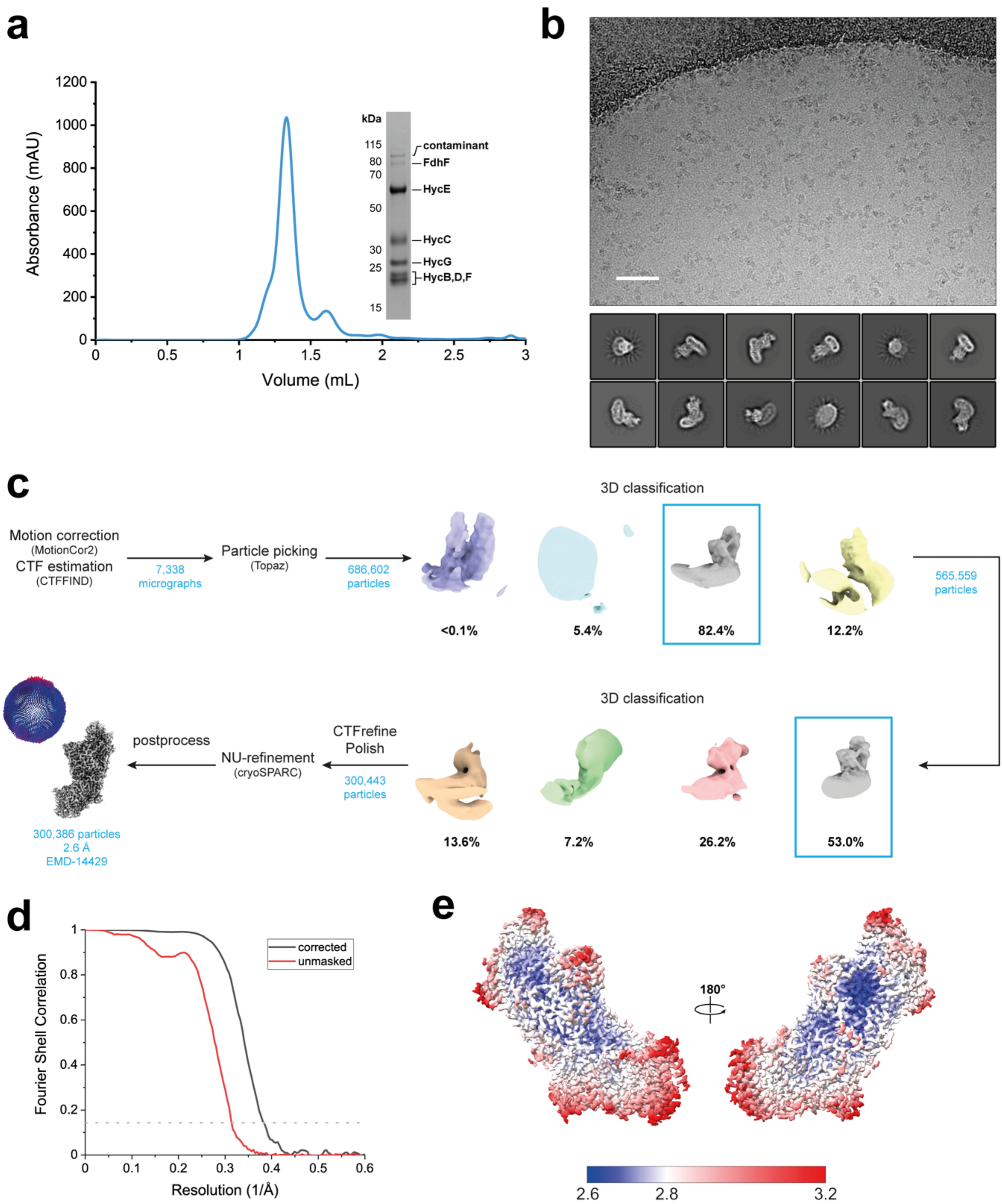


**Fig. S2 Sample preparation and data processing of anaerobically prepared FHL.** (**a**) Size exclusion chromatography (SEC) profile (Superdex 200 5/150 GL) and SDS-PAGE (right peak shoulder). Gel bands are labeled according to molecular mass. Traces of a ~80 kDa band could be identified in the gel, but no density for FdhF was observed in any 2D or 3D images of the cryo-EM dataset. A higher molecular weight band is a putative contaminant. (**b**) Representative cryo-EM micrograph (scale bar 500 Å) and 2D class averages. (**c**) Processing workflow, with all processing steps performed in RELION-3.1 except when otherwise indicated. The angular distribution plot was generated from the non-uniform refinement using pyem[1]. (**d**) Fourier Shell Correlation (FSC) curve of the final postprocessed map. (**e**) Local resolution estimation of the final postprocessed map.

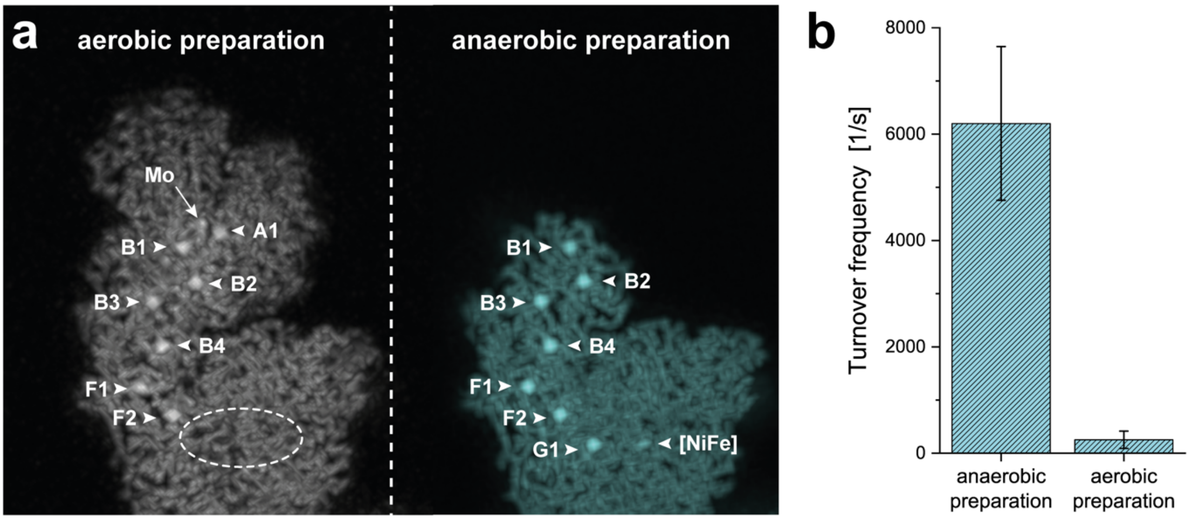


**Fig. S3 Comparison of aerobically and anaerobically prepared FHL** (**a**) Solid view representation of unsharpened maps of aerobically prepared FHL shows strong intensity for the [4Fe4S] clusters in FdhF (A1), HycB (B1-4) and HycF (F1-2). Density for the proximal [4Fe4S] cluster in HycG and the [NiFe] cofactor in HycE are weak (the approximate position is indicated with an ellipse). The anaerobically prepared FHL shows strong density for the proximal [4Fe4S] cluster (G1) and the [NiFe] cofactor. (**b**) Hydrogen uptake activity of anaerobically (6201 ± 1445 $s^{-1}$) and aerobically (253 ± 164 $s^{-1}$) prepared FHL. Initial slopes were calculated from 13 measurements from 2 biological replicates for the anaerobic preparation and 4 measurements for the aerobic preparation.

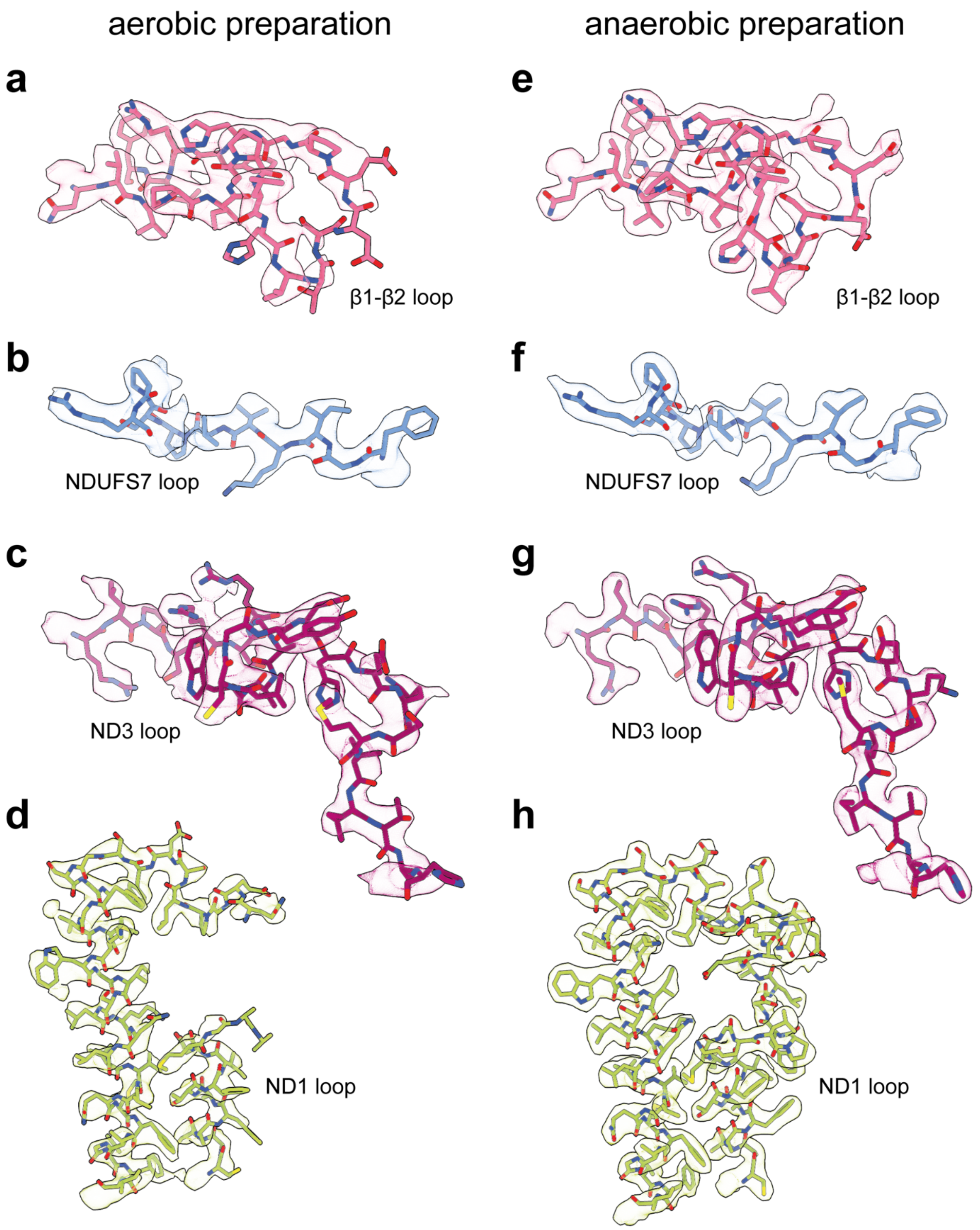


**Fig. S4 Comparison of aerobically and anaerobically-purified FHL in the loop region connecting the membrane and soluble arms.** Model and cryo-EM density for the conserved loop cluster in aerobically (**a-d**) and anaerobically (**e-h**) prepared FHL. The ND1 loop is partially unresolved for the aerobically prepared sample, whereas the anaerobically prepared sample shows side chain density for all loops.

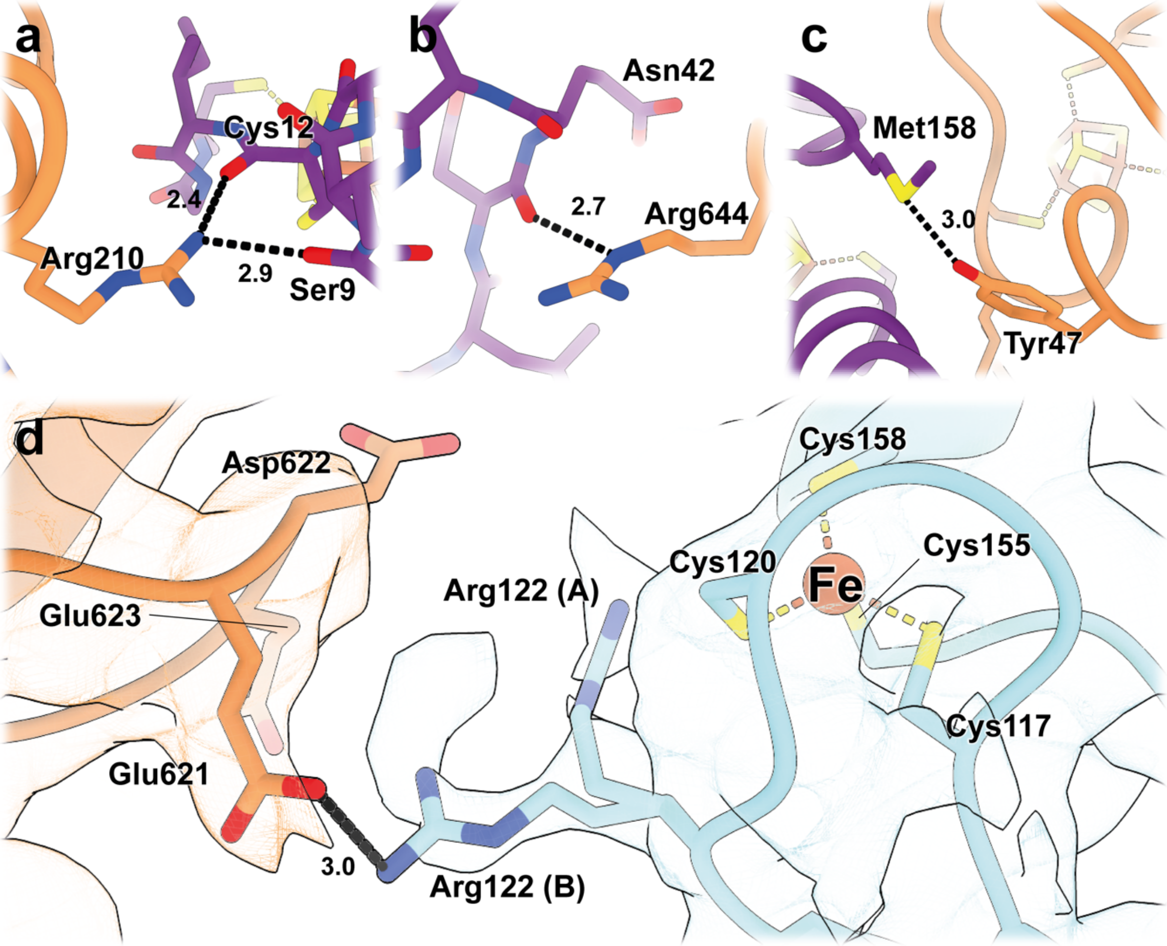


**Fig. S5 Interaction of the formate dehydrogenase subunit FdhF with HycB and HycF.** (**a-c**) Regions of hydrogen-bonding between FdhF and HycB (**d**) Interaction site between FdhF and HycF. Arginine-122 of HycF, which sits near to an unpredicted metal-ion of HycF and within a small electropositive region, shows two alternative sidechain conformations. In position A, it is close to the putative iron, while in position B, it forms a salt bridge with Glu621 of a negatively charged patch on FdhF (Fig. 2c).

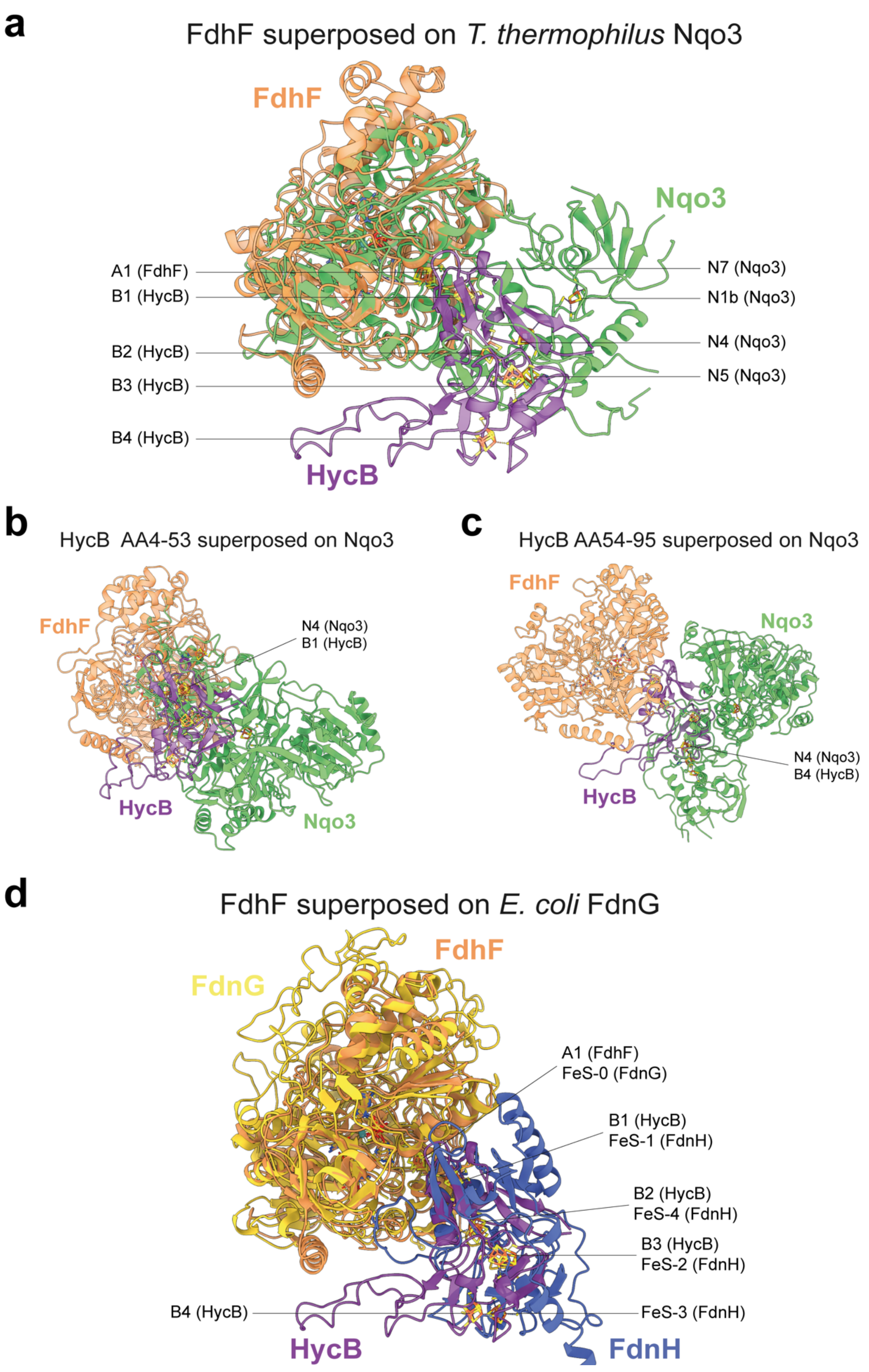


**Fig. S6 Superposition FdhF.** (**a**) *E. coli* FdhF can be superposed with the C-terminal region of *T. thermophilus* complex I Nqo3 (PDB 4HEA[2]) with an RMSD of 6.610 Å across all 496 atom pairs, bringing the [4Fe4S] cluster A1 in FdhF in register with the off-pathway [4Fe4S] cluster N7 in Nqo3. (**b**) HycB residues 4-53 can be superposed on the N-terminal region of Nqo3 with an RMSD of 10.306 Å over all 48 atom pairs, so that the [4Fe4S] cluster B1 (HycB) matches N4 (Nqo3), but the FdhF subunit does not match Nqo3. (**c**) HycB residues 54-95 can be superposed on the N-terminal region of Nqo3 with an RMSD of 14.609 Å across all 38 atom pairs, so that the [4Fe4S] cluster B4 (HycB) matches N4 (Nqo3), but the FdhF subunit does not match Nqo3. (**d**) FdhF superposed on the crystal structure of the *E. coli* formate dehydrogenase-N (PDB: 1KQF[3]) with an RMSD of 8.155 Å over 677 atom pairs, closely resembles the quarternary structure of the FdhF-HycB interaction. All [4Fe4S] clusters in FdhF and HycB of *E. coli* FHL match those of *E. coli* Fdh-N.

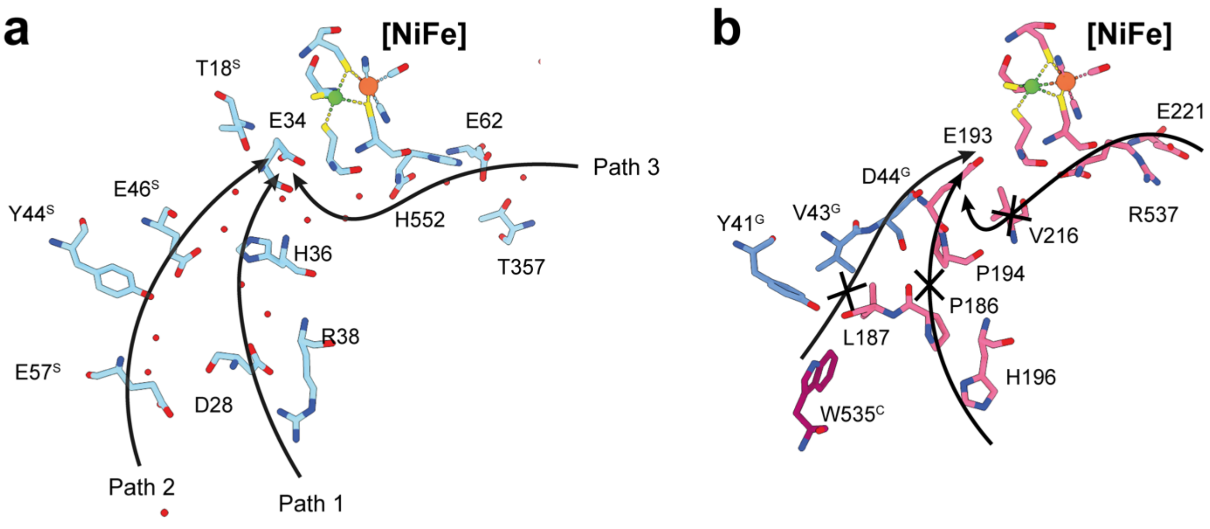


**Fig. S7 Substrate proton pathways in a soluble [NiFe] hydrogenase from *D. vulgaris* and FHL**. (**a**) Proton pathways observed in the crystal structure of *D. vulgaris* [NiFe] hydrogenase (PDB 4U9I[4]). Proton paths 1-3 leading to Glu34 as discussed in[4] are shown with arrows. (**b**) The residues forming these pathways are not conserved in *E. coli* FHL; in many cases they have been replaced by hydrophobic residues that appear to block proton transfer.

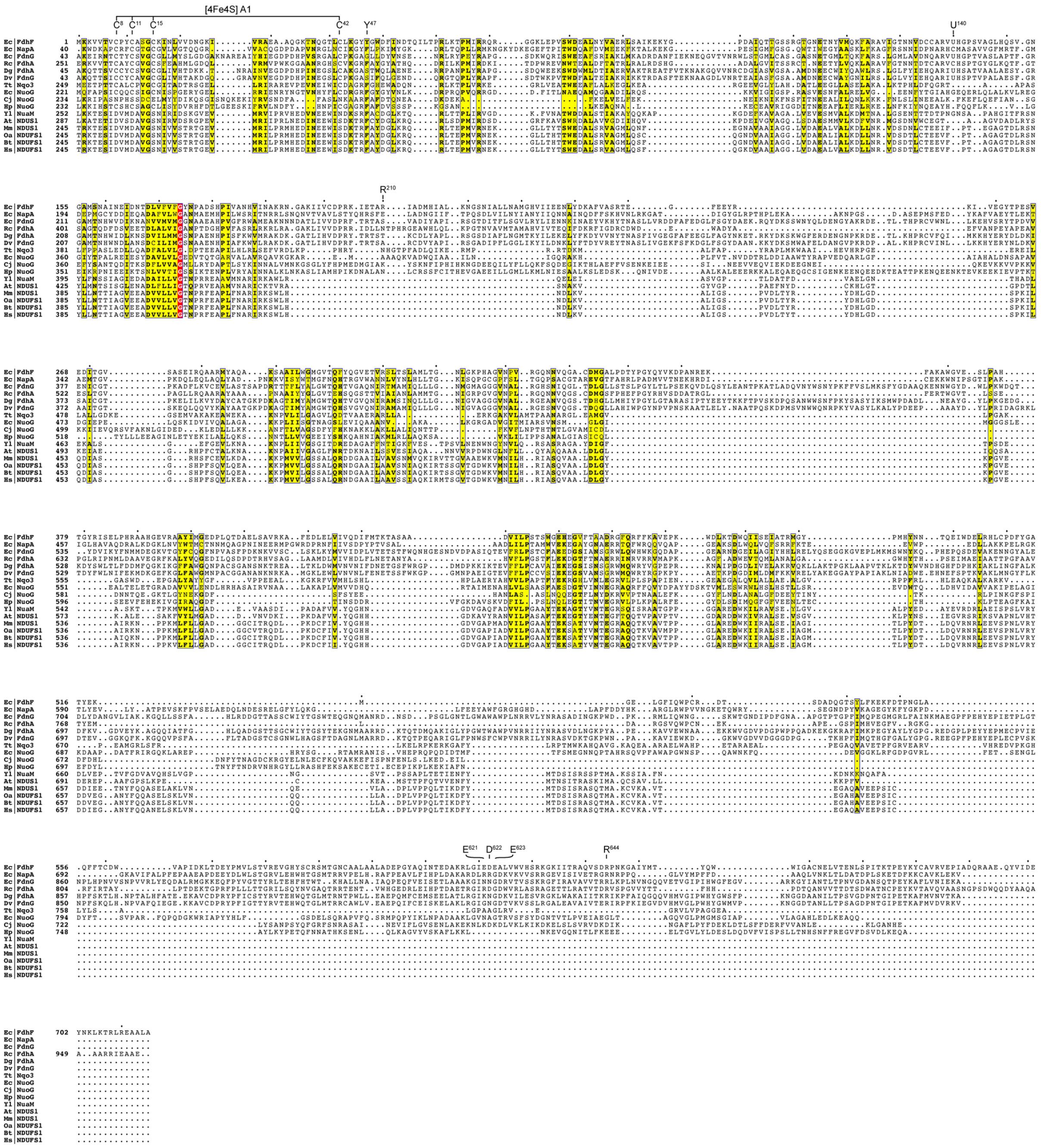


**Fig. S8 Sequence alignment of the formate dehydrogenase H (FdhF).** *Escherichia coli* formate dehydrogenase H (Ec|FdhF: P07658) was aligned with nitrate reductase from *E. coli* (Ec|NapA: P33937) and with formate dehydrogenases from *E. coli* (Ec|FdnG: P24183), *Rhodobacter capsulatus (*Rc|FdhA: D5AQH0), *Desulfovibrio gigas* (Dg|FdhA: Q934F5), and *Desulfovibrio vulgaris* (Dv|FdnG: Q72EJ1) as well as with the C-terminus of complex I subunit NDUFS1 from various species: *Thermus thermophilus* (Tt|Nqo3: Q56223), *E. coli* (Ec|NuoG: P33602), *Campylobacter jejuni* (Cj|NuoG: Q0P855), *Heliobacter pylori* (Hp|NuoG: I9X510), *Yarrowia lipolytica* (Yl|NuaM: Q9UUU3), *Arabidopsis thaliana* (At|NDUS1: Q9FGI6), *Mus musculus* (Mm|NDUFS1: Q91VD9), *Ovis aries* (Oa|NDUFS1: W5QB34), *Bos taurus* (Bt|NDUFS1: P15690), *Homo sapiens* (Hs|NDUFS1: P28331). The selenocysteine (U140) and cysteines coordinating cluster A1 in FdhF are marked (C8, C11, C15, C42). Residues interacting with HycB (Y47, R210, R544) and with HycF (E621, D622, E623) are marked.

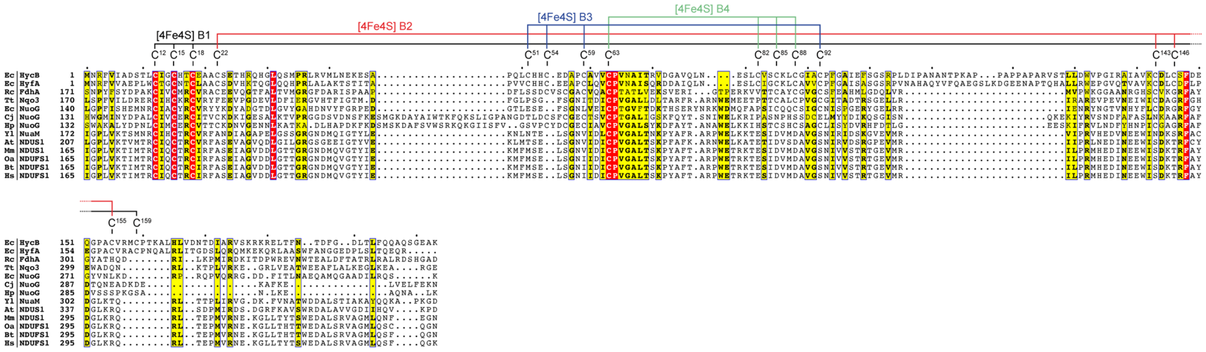

**Fig. S9 Sequence alignment HycB**. *Escherichia coli* HycB (Ec|HycB: P0AAK1) was aligned with homologue *E. coli* FHL-2 (Ec|HyfA: P23481), *Rhodobacter capsulatus* (Rc|FdhA: D5AQH0) and the N-terminus of complex I subunit NDUFS1 from various species: *Thermus thermophilus* (Tt|Nqo3: Q56223), *E. coli* (Ec|NuoG: P33602), *Campylobacter jejuni* (Cj|NuoG: Q0P855), *Heliobacter pylori* (Hp|NuoG: I9X510), *Yarrowia lipolytica* (Yl|NuaM: Q9UUU3), *Arabidopsis thaliana* (At|NDUFS1: Q9FGI6), *Mus musculus* (Mm|NDUFS1: Q91VD9), *Ovis aries* (Oa|NDUFS1: W5QB34), *Bos taurus* (Bt|NDUFS1: P15690), *Homo sapiens* (Hs|NDUFS1: P28331). Cysteines ligating cluster B1 (C12, C15, C18, C159), B2 (C22, C143, C146, C155), B3 (C51, C54, C59, C92) and B4 (C63, C82, C85, C88) are marked.

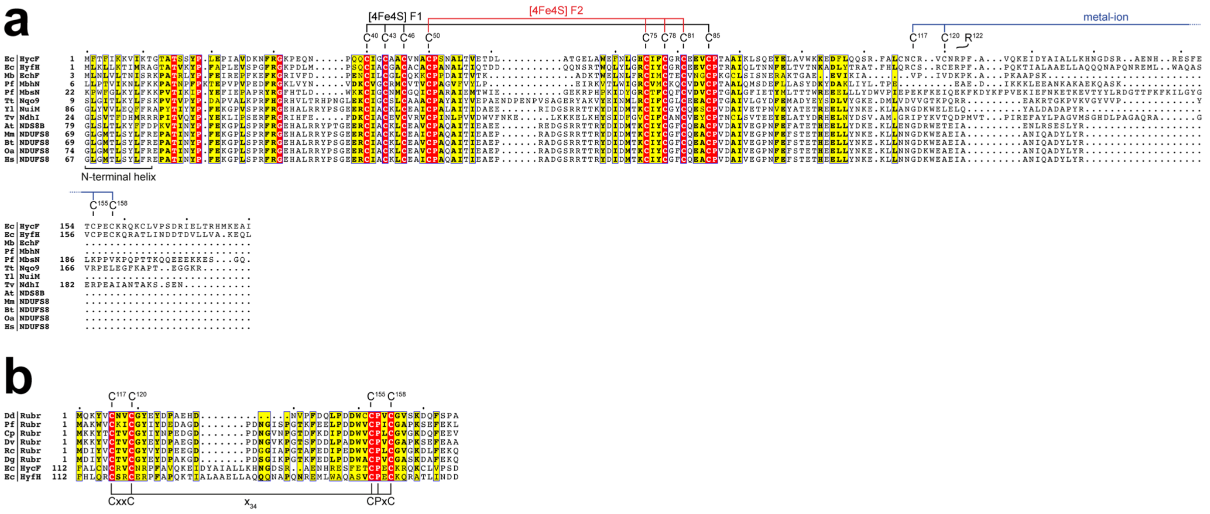


**Fig. S10 Sequence alignment HycF.** (**a**) *Escherichia coli* HycF (Ec|HycF: P16432) was aligned with its homologues in the complex I superfamily: *E. coli* (Ec|HyfH: P77423), *Methanosarcina barkeri* (Mb|EchF: O59657), *Pyrococcus furiosus* (Pf|MbhN: I6U851), *P. furiosus* (Pf|MbsN: I6U853), *Thermus thermophilus* (Tt|Nqo9: Q56224), *Yarrowia lipolytica* (Yl|NuiM: Q9UUT8), *Thermosynechococcus elongatus* (Tv|NdhI: Q8DL31), *Arabidopsis thaliana* (At|NDS8B: Q9FX83), *Mus musculus* (Mm|NDUFS8: Q8K3J1), *Bos taurus* (Bt|NDUFS8: P42028), *Ovis aries* (Oa|NDUFS8: A0A7M4DUG4), *Homo sapiens* (Hs|NDUFS8: O00217). Cysteines ligating cluster F1 (C40, C43, C46, C85) and cluster F2 (C50, C75, C78, C81) and cysteines coordinating the unpredicted metal ion (C117, C120, C155, C158) as well as R122, which interacts with FdhF are marked. The N-terminal helix of HycF is a common feature in the complex I superfamily. (**b**) *E. coli* (Ec|HycF: P16432) and (Ec|HyfH: P77423) was aligned with rubredoxins from various species: *Desulfovibrio desulfuricans* (Dd|Rubr: P04170), *Pyrococcus furiosus* (Pf|Rubr: P24297), *Clostridium pasteurianum* (Cp|Rubr: P00268), *Desulfovibrio vulgaris* (Dv|Rubr: P00269), *Ruminiclostridium cellulolyticum* (Rc|Rubr: Q9X709), *Desulfovibrio gigas* (Dg|Rubr: P00270). The unpredicted metal ion-binding motif (CxxC-$x_{34}$-CPxC) of HycF is marked.

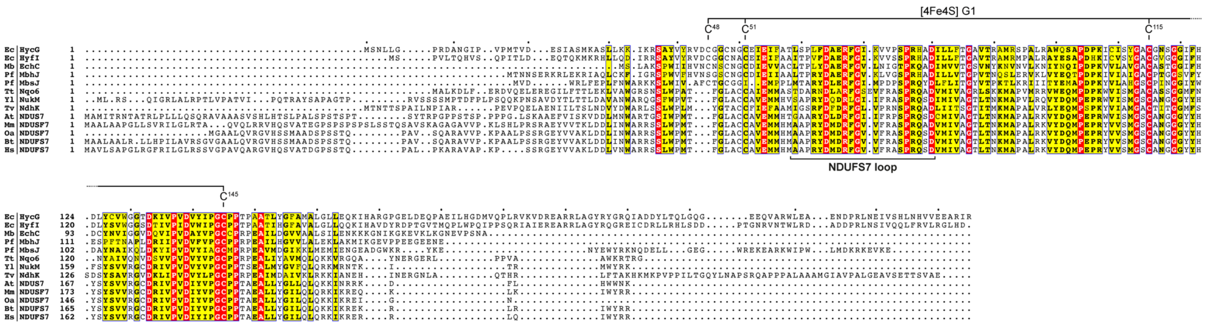

**Fig. S11 Sequence alignment HycG.** *Escherichia coli* HycG (Ec|HycG: P16433) was aligned with its homologues in the complex I superfamily: *E. coli* (Ec|HyfI: P77668), *Methanosarcina barkeri* (Mb|EchC: O59654), *Pyrococcus furiosus* (Pf|MbhJ: Q8U0Z8), *P. furiosus* (Pf|MbsJ: I6UZV3), *Thermus thermophilus* (Tt|Nqo6: Q56218), *Yarrowia lipolytica* (Yl|NukM: Q9UUT7), *Thermosynechococcus elongatus* (Tv|NdhK: Q8DKZ4), *Arabidopsis thaliana* (At|NDUS7: Q42577), *Mus musculus* (Mm|NDUSF7: Q9DC70), *Ovis aries* (Oa|NDUSF7: W5PPP6), *Bos taurus* (Bt|NDUFS7: P42026), *Homo sapiens* (Hs|NDUFS7: O75251). Cysteines ligating the proximal cluster G1 (C45, C51, C115, C145) and the NDUFS7 loop region are marked.

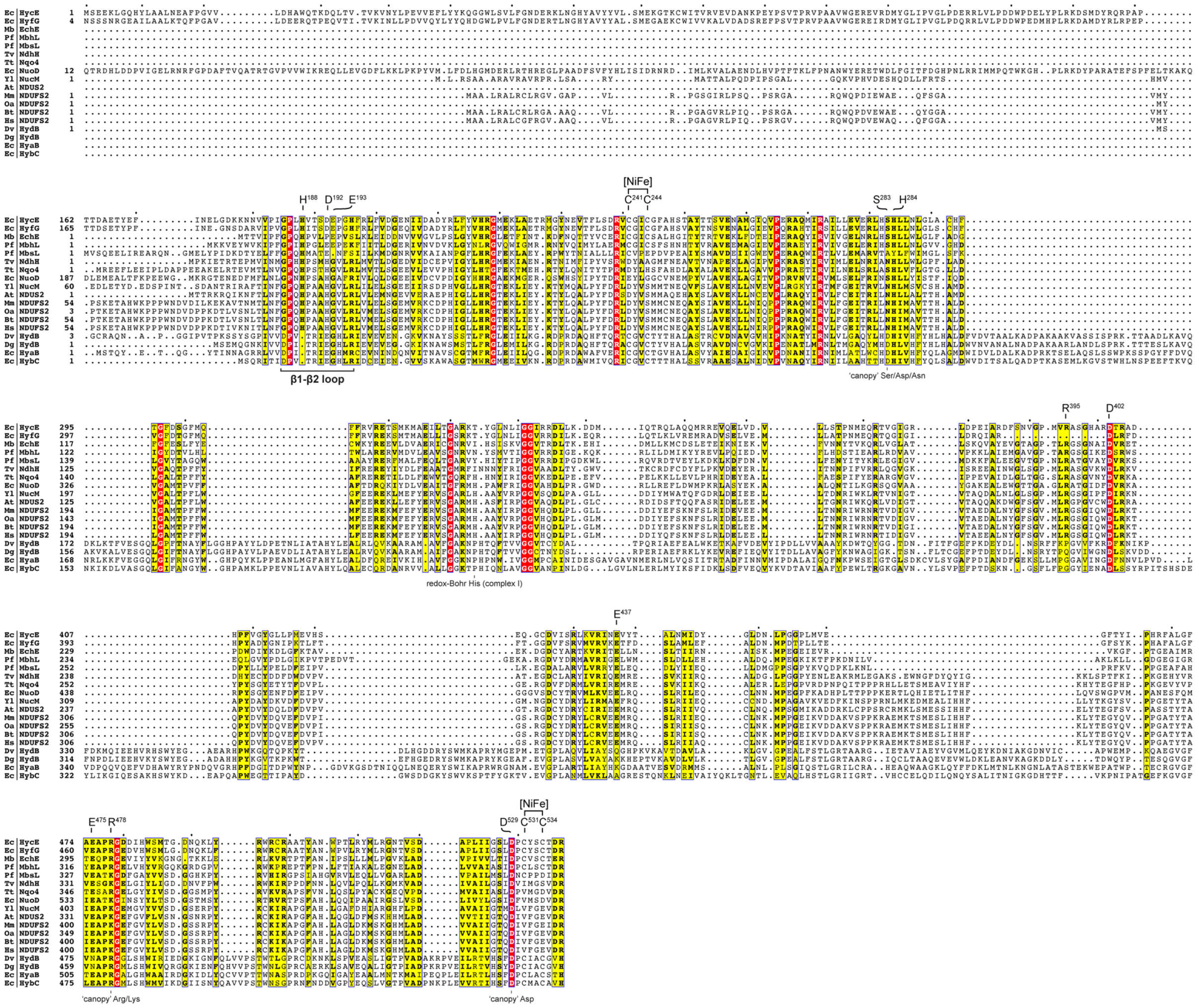


**Fig. S12 Sequence alignment HycE.** *Escherichia coli* (Ec|HycE: P16431) was aligned with its homologues in the complex I superfamily: *E. coli* (Ec|HyfG: P77329), *Methanosarcina barkeri* (Mb|EchE: O59656), *Pyrococcus furiosus* (Pf|MbhL: Q8U0Z6), *P. furiosus* (Pf|MbsL: I6V297), *Thermosynechococcus elongatus* (Tv|NdhH: Q8DJD9), *Thermus thermophilus* (Tt|Nqo4: Q56220), *E. coli* (Ec|NuoD: P33599), *Yarrowia lipolytica* (Yl|NucM: Q9UUU1), *Arabidopsis thaliana* (At|NDUS2: P93306), *Mus musculus* (Mm|NDUFS2: Q91WD5), *Ovis aries* (Oa|NDUFS2: W5PJ73), *Bos taurus* (Bt|NDUFS2: P17694), *Homo sapiens* (Hs|NDUFS2: O75306). And with soluble [NiFe] hydrogenases from *Desulfovibrio vulgaris* (Dv|HydB: P21852) and *Desulfovibrio gigas* (Dg|HydB: P12944) as well as *E. coli* Hyd-1 (Ec|HyaB: P0ACD8) and *E. coli* Hyd-2 (Ec|HybC: P0ACE0). Cysteines (C241, C244, C531, C534) are ligating the [NiFe] cofactor. The 'canopy' residues (S283, R478, D529), residues of the putative substrate proton pathway (H284, R395, D402, E437, E475) and residues (D192, E193) on the β1-β2 loop region are marked. The alignment represents the protein sequences after proteolytic maturation.

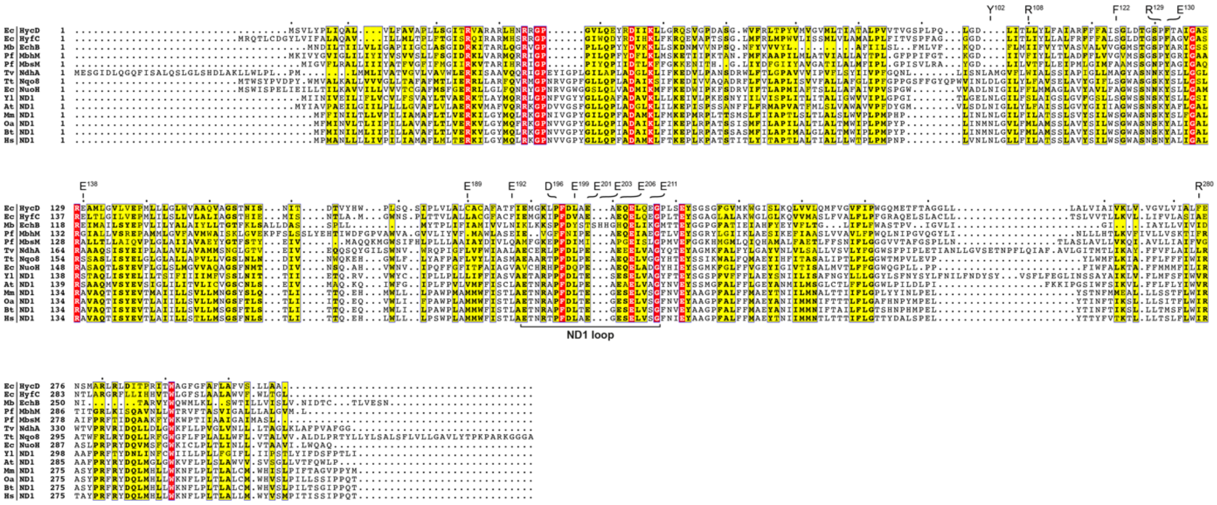


**Fig. S13 Sequence alignment HycD.** *Escherichia coli* (Ec|HycD: P16430 ) was aligned with its homologues in the complex I superfamily: *E. coli* (Ec|HyfC: P77858), *Methanosarcina barkeri* (Mb|EchB: O59653), *Pyrococcus furiosus* (Pf|MbhM: I6UQM0), *P. furiosus* (Pf|MbsM: I6V2A1), *Thermosynechococcus elongatus* (Tv|NdhA: Q8DL32),*Thermus thermophilus* (Tt|Nqo8: Q60019), *E. coli* (Ec|NuoH: P0AFD4), *Yarrowia lipolytica* (Yl|ND1: Q9B6E8), *Arabidopsis thaliana* (At|ND1: P92558), *Mus musculus* (Mm|ND1: P03888), *Ovis aries* (Oa|ND1: O78747), *Bos taurus* (Bt|ND1: P03887), *Homo sapiens* (Hs|ND1: P03886). Residues of the E-channel as well as the ND1 loop are marked.

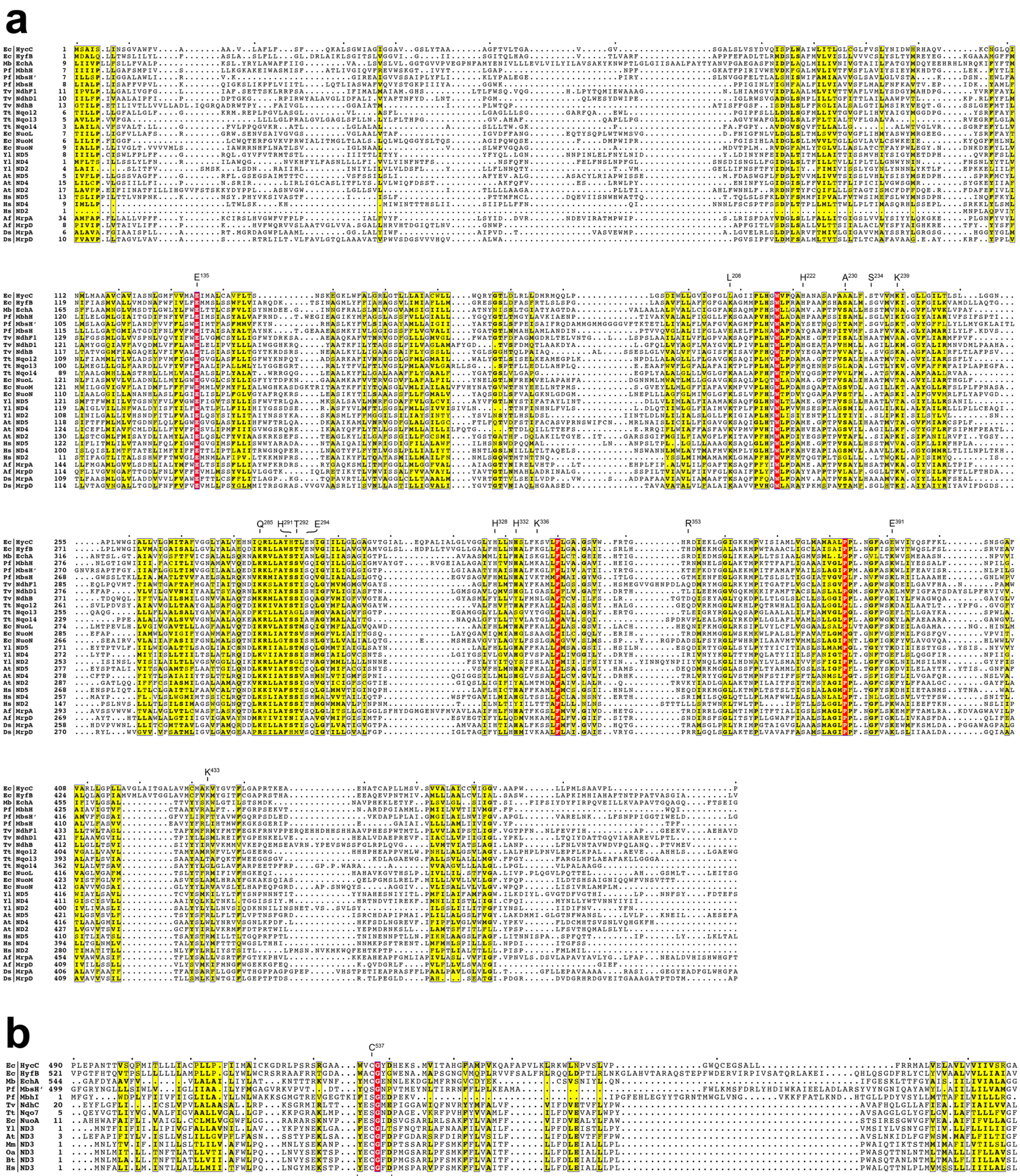


**Fig. S14 Sequence alignment HycC.** (**a**) Alignment of *Escherichia coli* HycC (Ec|HycC: P16429) residue 1-490 with antiporter-like subunits in the complex I superfamily: *E. coli* (Ec|HyfB: P23482), *Methanosarcina barkeri* (Mb|EchA: O59652), *Pyrococcus furiosus* (Pf|MbhH: I6UQL5, Pf|MbsH': I6TXQ1, Pf|MbsH: I6UZV7), *Thermosynechococcus elongatus* (Tv|NdhF1: Q8DKX9, Tv|NdhD1: Q8DKY0, Tv|NdhB: Q8DMR6), *Thermus thermophilus* (Tt|Nqo12: Q56227, Tt|Nqo13: Q56228, Tt|Nqo14: Q56229), *E. coli* (Ec|NuoL: P33607, Ec|NuoM: P0AFE8, Ec|NuoN: P0AFF0), *Yarrowia lipolytica* (Yl|ND5: Q9B6D3, Yl|ND4: Q9B6D6, Yl|ND2: Q9B6C8), *Arabidopsis thaliana* (At|ND5: P29388, At|ND4: P93313, At|ND2: O05000), *Homo sapiens* (Hs|ND5: P03915, Hs|ND4: P03905, Hs|ND2: P03891). And with antiporter-like subunits of MRP: *Anoxybacillus flavithermus* (Af|MrpA: B7GL84, Af|MrpD: B7GL98), *Dietzia sp.* (Ds|MrpA: A0A221C8X2, Ds|MrpD: A0A221C8X0). Residues referred to in Figure 5c are marked. (**b**) Alignment of *E. coli* HycC residues 490-608 with the C-terminal part of related antiporter-like subunits in the complex I superfamily and with complex I subunit ND3 from various species: *P. furiosus* (Pf|MbhI: I6U847), *T. elongatus* (Tv|NdhC: Q8DJ02), *T. thermophilus* (Tt|Nqo7: Q56217), *E. coli* (Ec|NuoA:

P0AFC3), *Y. lipolytica* (Yl|ND3: Q9B6C7), *A. thaliana* (At|ND3: P92533), *M. musculus* (Mm|ND3: P038993), *Ovis aries* (Oa|ND3: O78753), *Bos taurus* (Bt|ND3: P03898), *Homo sapiens* (Hs|ND3: P03897).
